# A separable neural code in monkey IT enables perfect CAPTCHA decoding

**DOI:** 10.1101/2020.04.12.038117

**Authors:** Harish Katti, S. P. Arun

## Abstract

Reading distorted letters is easy for us but so challenging for machine vision that it is used on websites as CAPTCHAs (Completely Automated Public Turing test to tell Computers and Humans Apart). How does our brain solve this problem? One solution is to have neurons invariant to letter distortions but selective for letter combinations. Another is for neurons to separately encode letter distortions and combinations. Here, we provide evidence for the latter using neural recordings in the monkey inferior temporal (IT) cortex. Neurons encoded letter distortions as a product of letter and distortion tuning, and letter combinations as a sum of letters. These rules were sufficient for perfect CAPTCHA decoding and were also present in neural networks trained for word recognition. Taken together, our findings suggest that a separable neural code enables efficient letter recognition.

## INTRODUCTION

Reading relies upon our ability to recognize letters and words despite variations in appearance. These include variations due to size, location or viewpoint, but also arising from writing styles, font and distortions of the writing surface. Yet we do solve this task so much better than computers that it is exploited on websites as CAPTCHAs (Completely Automated Public Turing test to tell Computers and Humans Apart) (von Ahn et al., 2003). How does the brain solve this problem? A widely held view is that neurons in higher visual areas are invariant to letter distortions and selective for specific combinations of letters (Grainger and Whitney, 2004; Dehaene et al., 2005). In this scenario, neural responses to letter combinations cannot be understood using single letters because of the presence of emergent features (Figure 1). This view is supported by observations that neurons in the monkey inferior temporal (IT) cortex, an area critical for recognition (Tanaka, 1996; DiCarlo et al., 2012), contain neurons invariant to size, location and viewpoint (Ito et al., 1995; Zoccolan et al., 2007; Ratan Murty and Arun, 2015), but selective for feature combinations (Kobatake and Tanaka, 1994; Baker et al., 2002; Brincat and Connor, 2004; Yamane et al., 2008). This selectivity for feature combinations increases with discrimination learning (Baker et al., 2002; Dehaene et al., 2005).

**Figure 1.**
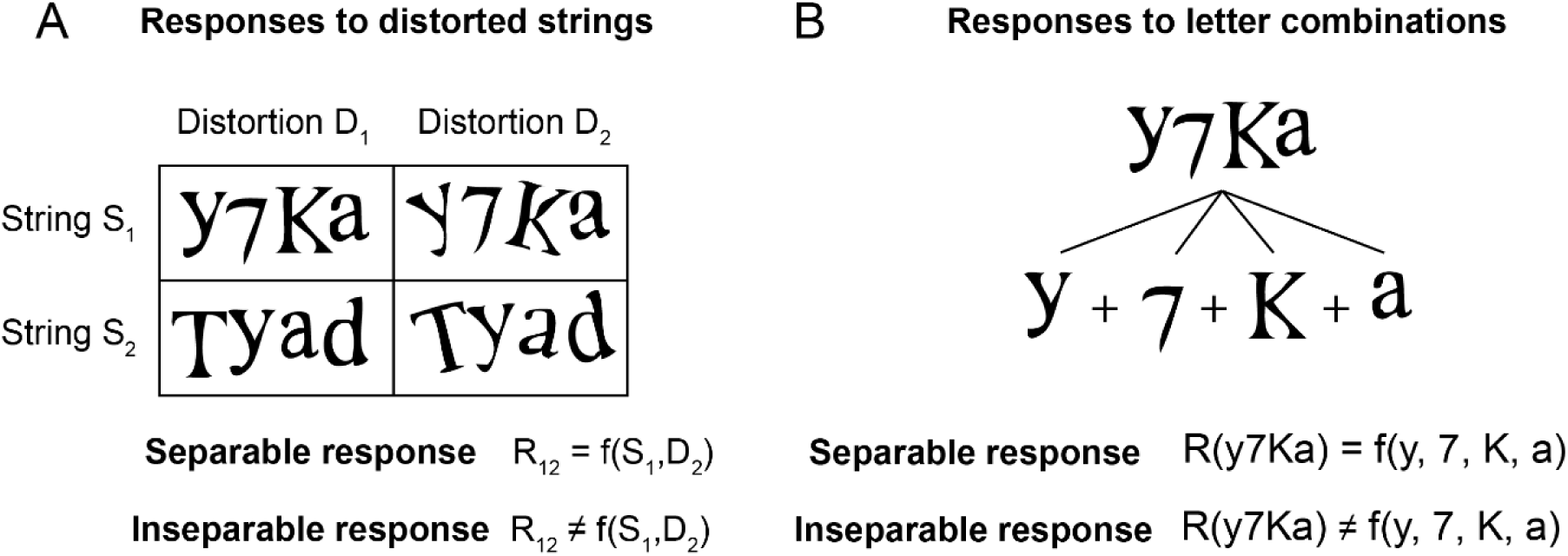
Separability of distortions and letter combinations. (A) We define the neural response to distorted strings as separable if the response to a particular string at a particular distortion can be understood as any function of string tuning and distortion tuning, and inseparable otherwise. Note that the underlying distortion tuning cannot be measured directly but can be estimated using a suitable model. (B) We define the neural response to letter combinations as separable if the response to a particular letter combination (e.g. y7Ka) can be understood as any function of the individual letter responses (e.g. y, 7, K and a separately), and inseparable otherwise. Inseparable responses could occur for instance if neurons were selective to specific combinations of letters. We note that responses to individual letters could either be measured directly or can be estimated by measuring responses to many letter strings containing the same letters.

An alternate possibility is that neurons may encode letters and their distortions as well as longer strings of letters in a separable fashion (Figure 1). By separable encoding we mean that responses to letter distortions or combinations can be understood in terms of their constituent letter or distortion selectivity. Evidence for this account comes from recent studies showing that IT neurons separably encode object identity and invariant attributes (Ratan Murty and Arun, 2018) and respond to objects as a sum of their parts (Zoccolan et al., 2005; Sripati and Olson, 2010; Pramod and Arun, 2018). This separability increases with reading expertise in humans (Agrawal et al., 2019).

However, these possibilities have not been investigated in IT neurons by recording their responses to distorted letter strings, which are unlike natural objects in that their constituent units (letters) are spatially separated and can undergo a wide variety of natural as well as stylistic distortions. Thus, it is not clear if the pre-existing neural machinery in naïve monkey IT itself suffices to solve CAPTCHAs.

To address this issue, we recorded neural responses in IT cortex of monkeys trained to fixate distorted letter strings but with no explicit training to discriminate or recognize these stimuli. We found that IT neurons encode distorted letter strings according to two systematic rules: letter distortions were encoded as a product of letter and distortion tuning, and letter combinations were encoded as a sum of single letters. To confirm that these rules are indeed sufficient to solve CAPTCHAs, we instantiated these rules in artificial neurons and asked whether these neural responses can be used to decode CAPTCHAs. Finally, we show that these rules naturally emerge in later layers of deep neural networks trained for letter and word recognition.

## RESULTS

We recorded from 139 single neurons in the IT cortex of two monkeys performing a fixation task. We tested each neuron with a fixed set of stimuli which comprised CAPTCHA-like strings that were 2-6 letters long, each of which was presented at four distortions (Figure S1). The stimuli also included single letters presented at each retinal location and at each distortion. This design enabled us to answer two fundamental questions regarding the separability of neural responses (Figure 1). First, can neural responses to letters and strings across distortions be predicted using their shape selectivity and distortion selectivity (Figure 1A)? Second, can neural responses to strings be explained using their responses to single letters (Figure 1B)?

### Do IT neurons show separability of letter shape, location and distortion?

We first characterized neural responses to single letters across retinal locations and distortions. The responses of an example IT neuron to each letter at every retinal location are shown in Figure 2A (the full set of responses to all stimuli can be seen in Figure S2). It can be seen that this neuron preserves its letter shape preference across retinal locations (Figure 2A, *top row*), and its location preference across letters (Figure 2A, *left column*) – suggesting that its response to letter and location is separable. Likewise, the response matrix for this neuron for each letter at every distortion is shown in Figure 2B – here too it can be seen that the neuron prefers some distortions over others and some letters over others. To quantitatively establish whether neural responses to shape x location x distortion combinations are separable, we compared two models in which the predicted response is either a sum or product of shape, location and distortion tuning. To avoid overfitting, we trained each model on odd-numbered repetitions, and evaluated its correlation on the even-numbered repetitions. We compared these model predictions with an estimate of neural reliability (split-half correlation; see Methods). The multiplicative model yielded excellent predictions of the neural response (*r* = 0.36, p < 0.0005; Figure 2C) that were close to the split-half correlation for this neuron (*r*_*sh*_ = 0.38, p < 0.0005). This correlation was even higher after removing the zero responses which were presumably because they were estimated from very few trials (r = 0.56, p < 0.0005). Thus, this neuron shows a response that is separable into shape, retinal location and distortion components.

**Figure 2.**
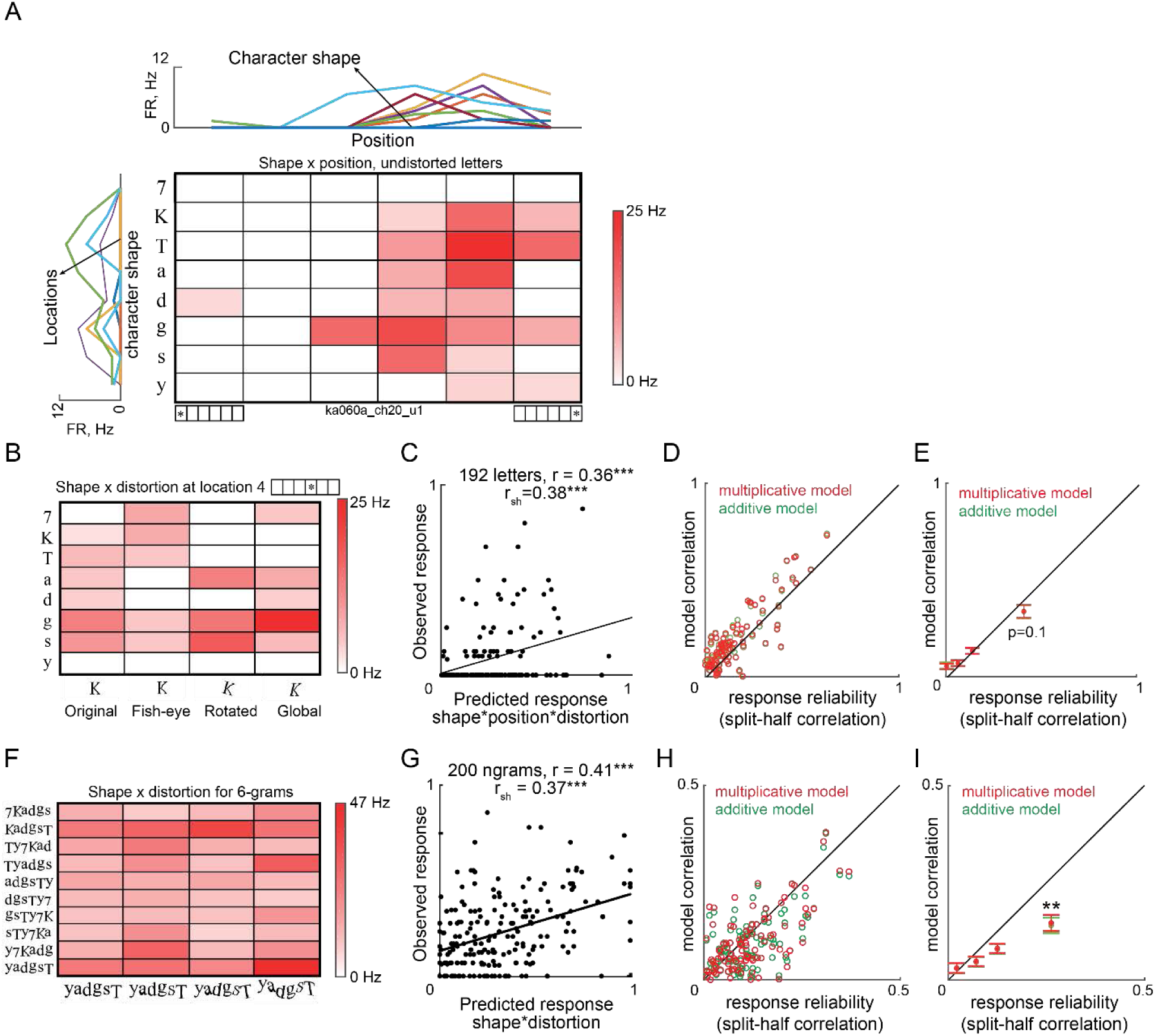
IT neurons show separable encoding of distorted letters and ngrams. (A) Colormap showing the firing rate of an example IT neuron to undistorted single characters at each retinal location. Darker colors indicate higher firing rate. *Above:* Responses across retinal locations for each character. *Left:* Responses to characters at each retinal location. (B) Colormap showing responses to each character at each of four distortions at a specific retinal location (location 4, 0.83° contralateral to fixation). (C) Observed responses (on even-numbered repetitions) to distorted letters across retinal locations plotted against predictions from the multiplicative model for the same neuron obtained from odd-numbered repetitions. Responses are normalized to the maximum firing rate. (D) Model correlation for the multiplicative (*red*) and additive (*green*) models plotted against response reliability across neurons with positive reliability. (E) Progression of model correlation with response reliability for each quartile of response variability for the multiplicative (*red*) and additive (*green*) models. Error bars represent standard error of the mean in each bin. The p-value indicates the outcome of a paired t-test comparing the additive and multiplicative model correlations for neurons belonging to that quartile. All quartiles showed no significant difference between the two models. (F) Colormap of responses to 6-grams at every distortion for the example neuron. (G) Observed responses to n-grams across distortions (from even-numbered repetitions) plotted against the multiplicative model predictions for the example neuron (trained on odd-numbered repetitions). Responses are normalized to the maximum firing rate. (H) Model correlation for the multiplicative (*red*) and additive (*green*) models plotted against response reliability across neurons with positive reliability. (I) Progression of model correlation with response reliability for each quartile of response variability for the multiplicative (*red*) and additive (*green*) models. Error bars represent standard error of the mean in each bin. Asterisks indicate statistical significance (** is p < 0.005) of a paired t-test comparing model correlations of the additive and multiplicative models for neurons belonging to that quartile. All other quartiles showed no significant difference.

Although this particular neuron shows separable responses, overall CAPTCHA decoding by the entire population might be driven by small numbers of neurons with highly consistent but inseparable responses. This would produce systematically worse model fits for neurons with high response consistency. We observed no such trend upon plotting model correlation against response reliability (Figure 2D). Rather, model correlations were close to the response reliability, indicating that they explained nearly all the explainable variance in the responses. We observed equivalent fits for the additive and multiplicative models for every level of response reliability (*r* = 0.053, 0.087, 0.12, 0.32, p < 0.05 for quartile-wise additive model fits and *r* = 0.05, 0.08, 0.12, 0.32, p < 0.05 for quartile-wise multiplicative model fits; Figure 2E).

### Do IT neurons show separability of strings and distortions?

Next we asked whether neural responses to longer strings across distortions are separable i.e. can be understood in terms of string and distortion selectivity. The responses of the same example neuron to 6-grams at multiple distortions is shown in Figure 2F. For each n-gram length (ranging from 2 to 6), we calculated the tuning for n-gram string as the average response to a given string (averaged across distortions) and the tuning for distortion as the response to a given distortion (averaged across strings) – and asked whether the response to n-gram strings across distortions can be understood in terms of the product or sum of tuning for n-gram string and tuning for distortion. The multiplicative model yielded excellent fits to the observed responses (r = 0.41, p < 0.0005; Figure 2G) that were comparable to the response reliability (*r*_*sh*_ = 0.37, p < 0.0005).

This pattern was true across the population: model fits were close to the response reliability for every level of reliability (Figure 2H). Furthermore, multiplicative models yielded slightly better fits than additive models (Figure 2H). This effect was most pronounced for the most reliable quartile of neurons (average correlations across 34 neurons with reliability > 0.13: r = 0.137 ± 0.1 for multiplicative model, r = 0.129 ± 0.1 for additive model; p = 0.005, paired t-test, Figure 2I).

We conclude that IT neurons show separable encoding of distorted letter strings.

### Do IT neurons show separability of responses to letter combinations?

Next we set out to characterize neural responses to letter combinations. The responses of an example IT neuron to n-grams varying in length from 2 to 6 letters is shown in Figure 3A. It can be seen that this neuron responded strongly to strings that contained the trigram “gsT” – but it also responds strongly to the letter “g” (Figure S2) suggesting that responses to longer strings can be explained using single letter responses. To quantitatively assess this possibility, we asked whether the response of each neuron to n-grams of each given length can be explained as a location-weighted sum of multiplicatively reconstructed responses to single letters (see Methods). The location-weighting accounts for the contribution of a given letter depending on its location within the full string. This model yielded excellent fits for the example neuron (r = 0.41, p < 0.0005; Figure 3B) that were equivalent to the response reliability (*r*_*sh*_ = 0.38, p < 0.0005). This pattern of results were true for the entire population: model fits approached or exceeded response reliability across neurons for every level of reliability (Figure 3C).

**Figure 3:**
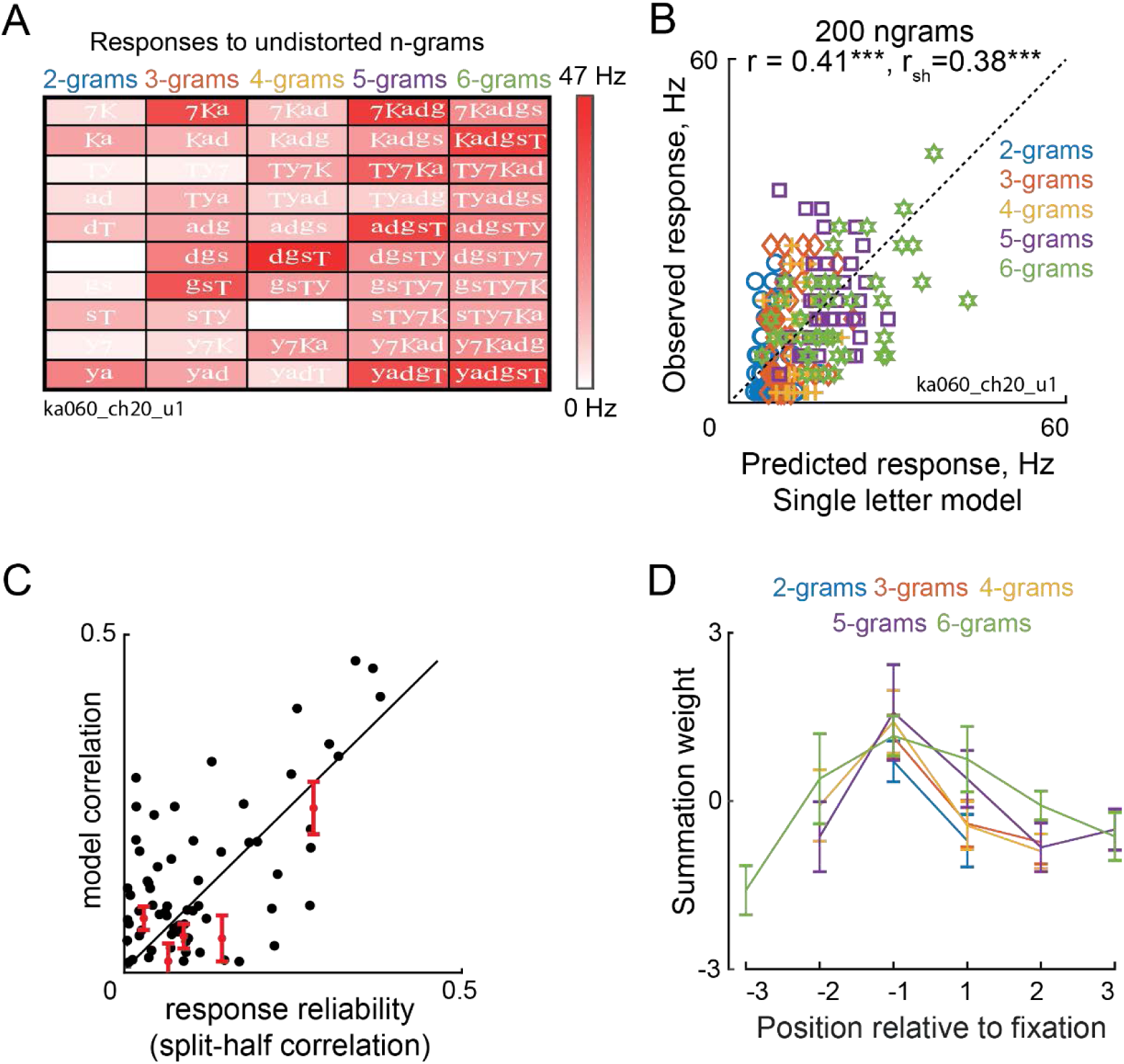
Neural responses to n-grams are compositional. (A) Response of the example neuron (same as Figure 1) to all undistorted n-grams in the experiment. (B) Observed responses to all n-grams (on even-numbered repetitions) plotted against the predictions of the single letter model (trained on odd-numbered repetitions) for the example neuron. (C) Model correlation for each neuron plotted against the response reliability across neurons. Average model correlations for each 20^th^ percentile of neurons with positive response reliability are shown in red, with error bars depicting s.e.m across each such group. (D) Average summation weights for each retinal location for different n-gram lengths, averaged across both cross-validated estimates (10-fold) and across neurons. Error bars depict s.e.m.

The above analysis shows that neural responses to letter strings is well approximated by a sum of single letter responses, but neurons might still show systematic model errors indicative of selectivity for letter combinations. To explore this possibility, we asked whether bigrams that have high model error show systematically larger errors for longer strings that contain these bigrams. This revealed no such systematic difference (Figure S3).

Next we examined the estimated summation weights at each retinal location. For each neuron, we normalized its response to the maximum firing rate across all stimuli, and estimated summation weights for each retinal location and each n-gram length (across all n-grams). The average summation weights are shown in Figure 3D. For all n-gram lengths, we observed larger weights of letters close to fixation compared to more peripheral locations. This is consistent with the well-known bias of IT neuron receptive fields towards fixation (Op De Beeck and Vogels, 2000). Moreover, the relationship between n-gram responses and the sum of single letter responses followed a characteristic 1/n progression (Figure S4). This is consistent with the divisive normalization observed previously in IT neurons (Zoccolan et al., 2005).

We conclude that IT neurons show separable encoding of letter strings whereby the responses to strings can be predicted using a sum of single letter responses.

### Is separability sufficient for CAPTCHA decoding?

The above analyses show that IT neurons encode distorted letter strings as a product of shape and distortion tuning, and encode letter combinations as a sum of single letters. However they do not establish that CAPTCHAs can be solved using these responses, or that they can be solved if responses are separable.

To address these issues, we first asked whether CAPTCHAs can be solved using the responses of IT neurons to the distorted letter strings. To this end, we trained a 9-way linear classifier to decode character identity at every retinal location (see Methods). Since each retinal location could have one of 8 possible letters or be a blank space, chance performance for this decoding scheme is 1/9 or 11%. We used 5-fold cross-validation to avoid overfitting. As expected, decoding accuracy using the recorded population of neurons was well above chance (average accuracy across 6 retinal locations = 29%).

The above result shows that we can decode letter identities successfully using a small population of neurons, but it is not clear how this decoding accuracy would scale with a larger population of neurons. It is also not clear whether the decoding accuracy is being driven by neurons with separable or inseparable responses. To resolve this issue, we asked whether CAPTCHAs can be decoded by a population of synthetic neurons with separable encoding.

We created 1,000 synthetic neurons that had either broad or sharp tuning for shape, location and distortion (see Methods; Figure 4A). We calculated neural responses to shape-location-distortion combinations either using an additive or multiplicative function. Their responses to longer strings was taken simply as a sum of their responses to single letters, and finally we included divisive normalization whereby the net response to longer strings is divided by the string length. To provide a tough challenge of the CAPTCHA solving ability of this neural population, we created a large set of n-grams (n = 17,920) of varying string lengths, which could even contain spaces between letters (see Methods). We then asked whether these separable neural responses could be used to decode character identity at each location as before. In this manner, we created several groups of neurons: broad tuning with multiplicative/additive integration with/without normalization, sharp tuning with multiplicative/additive integration with/without normalization and mixed tuning with multiplicative/additive integration with/without normalization.

**Figure 4:**
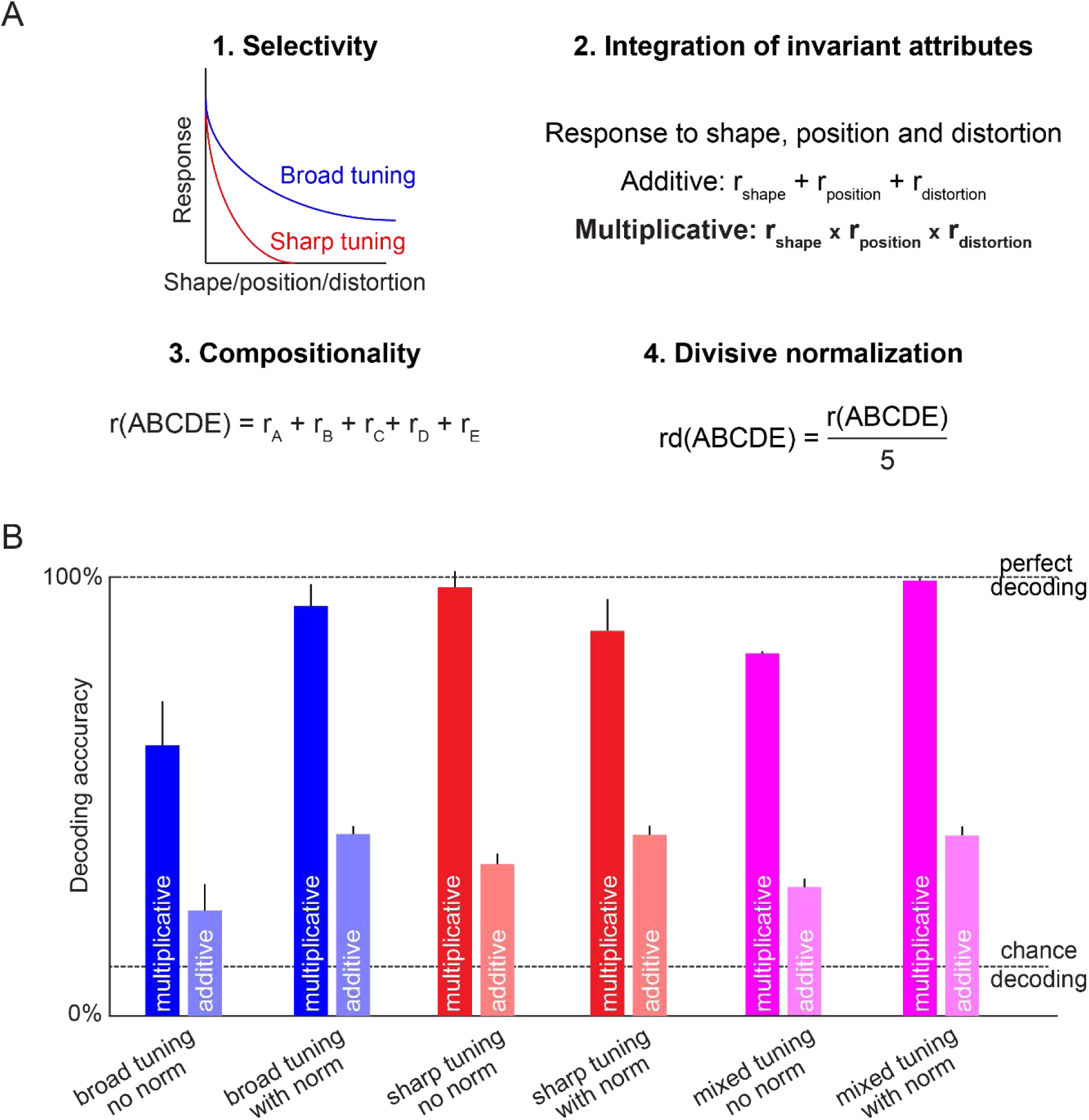
Response separability is sufficient for solving CAPTCHAs. (A) To explore what properties present in a population of separable neurons can solve CAPTCHAs, we first created 1,000 artificial neurons with either sharp or broad tuning for letter, location and distortion. Next, we calculated responses to distorted letters using either additive or multiplicative combination of shape, location and distortion tuning. To calculate responses to letter combinations, we simply added the responses to all the constituent letters. Finally, responses were (or were not) subjected to divisive normalization. We tested the CAPTCHA solving ability of these artificial neurons on a large set of 17,920 n-grams. (B) To evaluate this neural population for CAPTCHA decoding, we trained a linear classifier to decode which of 9 possible characters (8 letters + 1 blank) were present at each retinal location in each n-gram. The resulting average character decoding accuracy is shown for each neural population created with a specific set of choices. Error bars represent standard deviation over 100 iterations of sampling with replacement from a population of 1000 synthetic neurons.

The resulting character decoding accuracy for each of these artificial neuron populations is summarized in Figure 4B. We observed several interesting patterns. First, decoding accuracy was always better when shape and distortion tuning were multiplicatively combined. This is because multiplying two signals enables efficient decoding compared to adding them, as we have reported previously (Ratan Murty and Arun, 2018). Second, decoding was better when responses of neurons with broad or mixed tuning were divisively normalized than when they were not. This is because normalization avoids excessively large responses to increasing content in the visual field. Since n-grams of different lengths can have the same character at a given location (e.g.; “7yKa “and “Ty7Kad” both have “y” in the second location), longer n-grams will automatically have higher firing rates in the absence of divisive normalisation and will result in two different read-outs for the same character label. Third, decoding is worse when neurons with sharp tuning are subjected to divisive normalization, which happens because sharply tuned neurons respond to only a few retinal locations and divisive normalization results in excessive scaling by the total string length. Finally, and most importantly, we obtained near-perfect CAPTCHA decoding accuracy with a mixed population of neurons with sharp and broad tuning.

We conclude that response separability is sufficient to solve CAPTCHAs.

### Do neural networks trained for letter recognition show separability?

The above sections demonstrate that IT neurons show separable encoding of letter distortions and combinations. We then asked whether this is indeed an essential property of a visual system that aims to decode words and letters and is not merely restricted to biological neurons. Would separability be equally important even when the visual system was explicitly trained to identify entire words or n-grams as opposed to individual characters?

To address this issue, we analysed the responses of four deep convolutional networks. The first network served as a baseline since it was trained on a standard object classification task and not for letter recognition (denoted as *imagenet*) (Simonyan and Zisserman, 2014). The remaining three of these were trained on a large-scale annotated dataset of about 8 million images generated from strings of variable lengths and different choices of fonts, projective transformations and backgrounds (Jaderberg et al., 2014). The networks were trained with three different goals: (1) identifying the character at each of the 23 possible locations in the input word (denoted as *charnet*); (2) enumerating all n-grams contained within the input word (denoted as *ngramnet*) and (3) whole word identification from a dictionary of 90,000 words (denoted as *dictnet*). In all networks, we selected the penultimate fully connected layer (fc7, with 4096 units) for further analysis, because this layer has the same dimensionality and is subsequently used by the decision layer.

We next analysed the activations of each network to the stimuli used in the neural recordings. As before we fit additive and multiplicative models to predict the response of each unit in the fc7 layer to single letters across retinal locations and distortions. In all four networks, unit activations were better fit by the multiplicative model (Figure 5A). Interestingly, the *imagenet* network yielded lower model fits compared to the networks trained on word recognition, whereas the *dictnet* model fits were the highest among all networks. We obtained similar results on analysing n-gram responses across distortions (Figure 5A). Next we asked whether unit activations to longer n-gram strings in our dataset, can be explained using single letters. Here too we observed a clear progression, whereby the *imagenet* units yielded the least separability, and *dictnet* units were most separable (Figure 5B). However in all cases, model correlations are well below 1, suggesting that neural responses are not fully separable despite training. Such emergent compositionality has been observed in high-performing deep networks trained for natural scene categorisation (Zhou et al., 2014).

**Figure 5:**
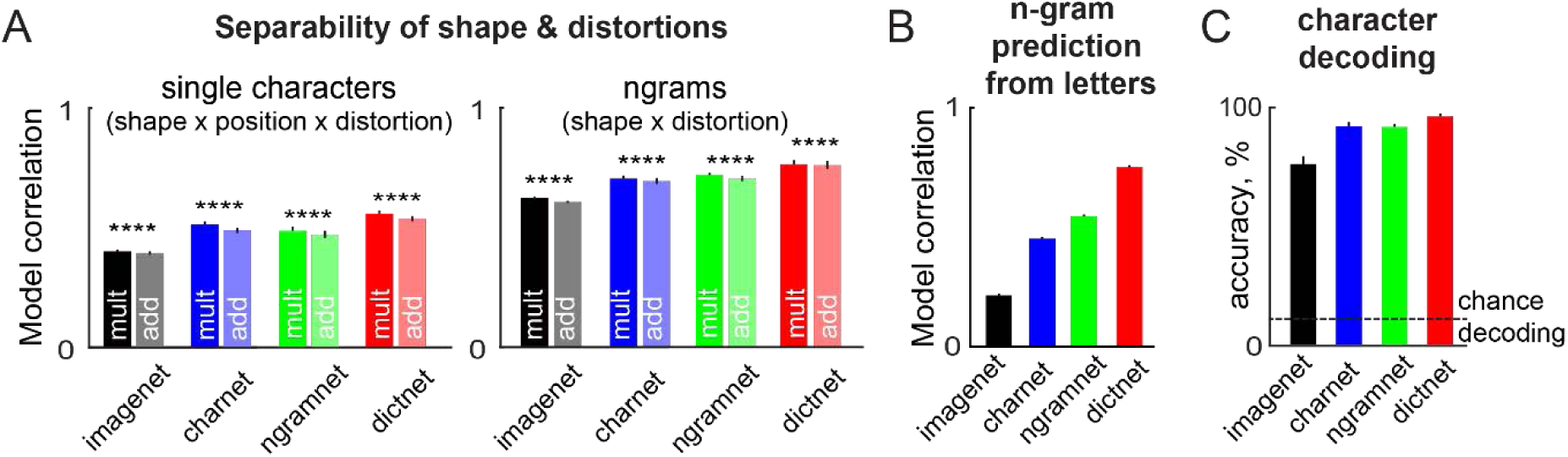
Compositionality in CNNs trained for text decoding. (A) *Left:* Model correlations for additive and multiplicative models for shape x location x distortion for each neural network. *Right:* Model correlations for additive and multiplicative models for shape x distortion for each neural network. Error bars represent s.e.m across neurons. Asterisks represent statistical significance calculated using a paired t-test on the additive and multiplicative model correlations across all 4096 units (**** is p < 0.00005). (B) Average model correlations for the linear model that predicts n-gram responses as a weighted sum of single letter responses. Error bars represent s.e.m. across neurons. (C) Character decoding accuracy across the stimulus set used in the neural recording experiment for each neural network. Error bars represent s.e.m over character decoding accuracy over 6 retinal locations.

Since the network trained on complete word recognition (*dictnet*) shows the greatest separability, we wondered whether its performance on letter recognition would also be the best across all networks. To investigate this issue, we compared the performance of each network on 9-way decoding of character decoding at each retinal location as before on the responses to our original n-gram set (50 unique n-grams x 4 distortions). As expected, the performance of *dictnet* was better than the other networks (decoding accuracy: 74%, 92%, 93% and 95% for *imagenet, charnet, ngramnet* and *dictnet*; Figure 5C).

Thus, training deep neural networks for word recognition leads to separable neural representations for distortions and letter combinations. We speculate further that imposing separability as a constraint during learning can lead to better character decoding performance.

### Can IT neurons decode both easy and hard CAPTCHAs?

The above results were based on testing IT neurons with distorted letter strings with spatially separate letters and relatively simple distortions. However we also included a small set of more challenging CAPTCHAs which contain overlapping letters or cluttered backgrounds (Figure 6A). We predicted that letter decoding should also be possible in these hard CAPTCHAs, albeit at a weaker level.

**Figure 6.**
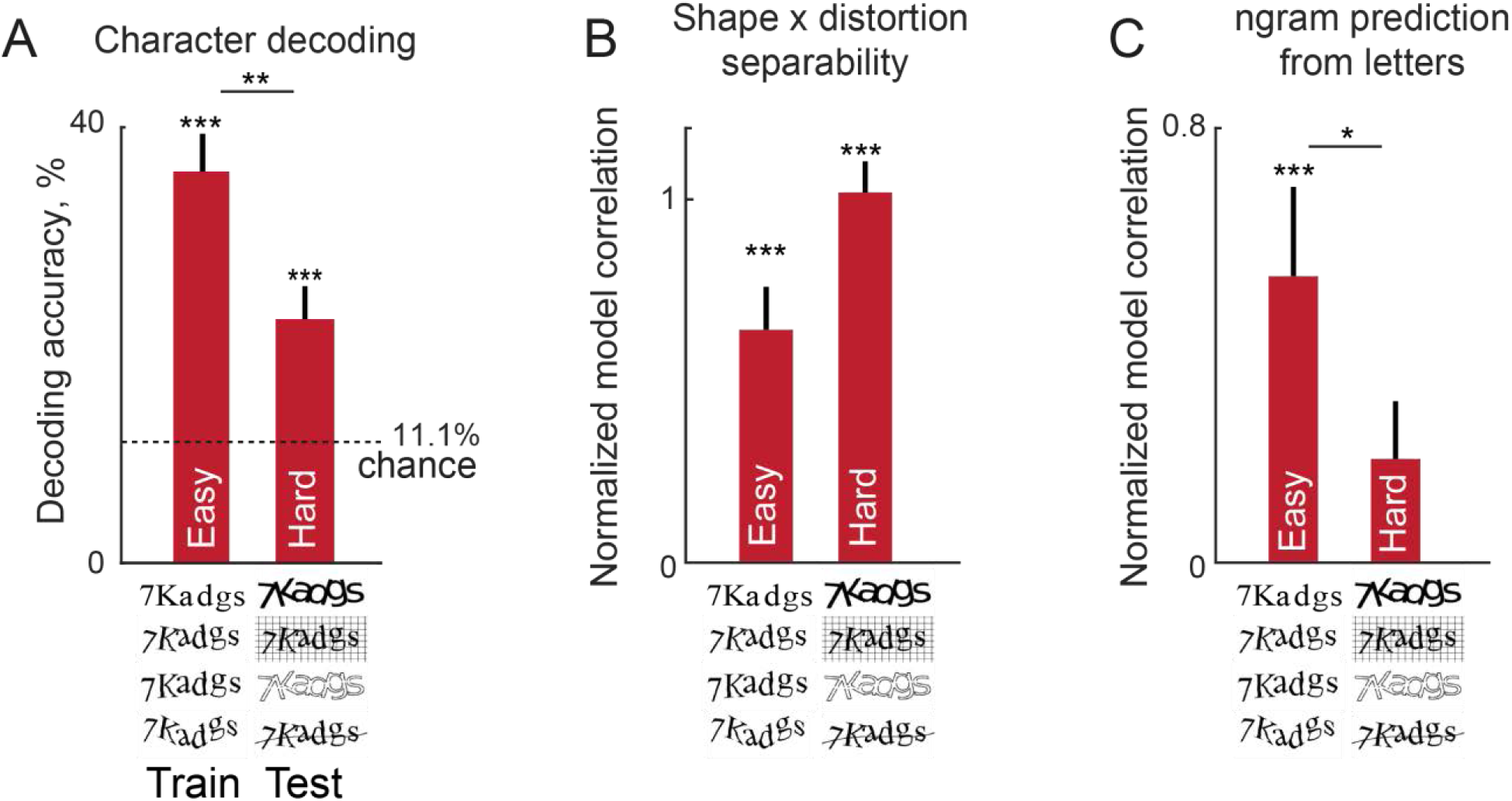
Decoding and separability for easy and hard CAPTCHAs by IT neurons. A) Average decoding accuracy (of 9 characters across all retinal locations) for classifiers trained on the easy CAPTCHAs (left) and tested on the hard CAPTCHA strings (right). Error bars represent s.e.m across decoding accuracy across all n-grams. Asterisks above each bar represent the statistical significance of the decoding accuracy compared to chance decoding which is 1/9 = 11% (*** is p < 0.0005, based on calculating the binomial probability of exceeding this number of correct labels out of a total of 8 strings x 6 retinal locations x 4 distortions = 192 responses given a binomial distribution with N = 192, p = 1/9 = 11.1%). Asterisks straddling the two bars represent the statistical significance of the difference in decoding accuracy (** is p < 0.005, rank-sum test for medians). B) Normalized model correlation for the shape x distortion multiplicative model, for the easy and hard CAPTCHA strings. Error bars represent s.e.m across neurons. For each neuron, we calculated the correlation between observed responses on even-numbered repetitions and the model predictions based on odd-numbered repetitions, and divided by the split-half correlation to obtain a normalized measure. Asterisks above each bar indicate whether the median normalized correlation is different from zero (*** is p < 0.0005, based on a sign-rank test across 139 neurons). Although the hard CAPTCHA strings show greater separability, this difference was not statistically significant (p = 0.2, signed-rank test across 139 neurons). C) Normalized model correlations for the n-gram responses predicted by single letter responses. Normalized correlations for hard CAPTCHA strings were not significantly different from zero (p = 0.09, signed-rank test), and this separability was significantly smaller than for the easy CAPTCHA strings (* is p < 0.05, signed-rank test across 139 neurons). All other conventions are as in B.

To assess this possibility, we took linear classifiers to perform 9-way character decoding at each retinal location as before, and asked whether they could decode letter identity from the hard CAPTCHA strings. Decoding accuracy was weaker for the hard CAPTCHAs but still remained significantly above chance (Figure 6A). Thus, CAPTCHAs that are hard for computer algorithms are also challenging for IT neurons.

Since hard CAPTCHAs are harder to decode, we wondered whether their responses are less separable with respect to distortions and string responses. To investigate this issue, we took the 8 six-letter strings that were common to both the easy and hard CAPTCHAs (with 4 distortions each), and asked whether these responses were separable in terms of distortions and n-gram responses. We first fit a shape x distortion multiplicative model to the responses to the 8 grams across 4 distortions. To compare the model performance, we divided the model correlation of each cell by its split-half correlation for each set of stimuli. Both easy and hard CAPTCHAs were equally separable (Figure 6B). Next we asked whether the average response of each n-gram (on even-numbered repetitions) can be predicted using a weighted sum of the single letter responses (with weights learned from responses on odd-numbered repetitions), and normalized the model correlation as before. The resulting normalized model correlations are shown in Figure 6C. As predicted, model correlations for hard CAPTCHAs were significantly smaller than for easy CAPTCHAs.

We conclude that hard CAPTCHAs are less separable than easy CAPTCHAs, which explains why they are harder to decode by the IT neural population.

## GENERAL DISCUSSION

Here we characterized the neural representation of distorted letters in IT neurons. We found two systematic encoding rules: neural responses to distorted letters was a product of distortion and shape tuning, whereas responses to letter combinations was a sum of single letter responses. These two rules were sufficient for perfect CAPTCHA solving and emerged in neural networks trained for word recognition. Below we discuss these findings in relation to the literature.

Our main finding is that there is a systematic separable code for distorted letter strings in monkey IT neurons, consisting of two rules. The first rule was that neural responses to distorted letters is approximated well by a product of distortion and shape tuning. This is consistent with recent observations that object identity and invariant attributes combine multiplicatively in IT neurons (Ratan Murty and Arun, 2018). By contrast, we have found the neural response to multiple objects to be a weighted sum of the constituent letters. This is consistent with several previous studies showing linear summation in IT neurons for objects (Zoccolan et al., 2005) as well as their parts (Sripati and Olson, 2010; Pramod and Arun, 2018). We speculate that IT neurons switch between multiplicative and additive integration of features depending on feature similarity.

Our findings are inconsistent with many previous studies suggesting that IT neurons are selective for specific combinations of features (Kobatake and Tanaka, 1994; Brincat and Connor, 2004; Yamane et al., 2008). However, these studies did not systematically compare object responses to their constituent parts or features. Doing this is nontrivial because isolating the parts from an object introduces new features where the object is cut – therefore a different approach is needed whereby the part responses can be estimated indirectly from the whole object responses. When this has been done, whole object responses were indeed predictable from the constituent parts, both in IT neurons (Sripati and Olson, 2010; Pramod and Arun, 2018) as well as in perception using visual search (Pramod and Arun, 2014, 2016, 2018).

Our findings are also inconsistent with a recent study showing that n-gram responses are not entirely predictable from single letters in IT neurons (Rajalingham et al., 2019). We speculate that the lower separability in that study could be due to more posterior recording sites in IT (posterior/central IT compared to ours which were in anterior ventral IT), different response noise distributions (multi-unit activity in that study vs single units in ours), and shorter presentation times (100 ms in their study with no inter-stimulus interval vs 200 ms on/off duration in ours), apart from of course stimulus and individual differences. These differences will need to be reconciled. However we have gone beyond these specific results to show that separability in neural responses are sufficient to solve CAPTCHAs. This of course does not imply that inseparable neural responses, with neurons tuned for letter combinations, cannot solve CAPTCHAs or are not useful for other purposes; such codes might be required for further downstream processing, such as binding combinations of letters into syllables (Dehaene et al., 2005, 2015). Rather, our results imply that, at least for the purposes of character decoding, response separability is sufficient. We speculate that there is a gradual progression in separability as well as invariance along the ventral pathway in order to enable efficient object recognition (DiCarlo et al., 2012).

Our results have important implications for how exactly letter and word representations change with reading. First, we note that perfect CAPTCHA decoding using a perfectly separable neural population (Figure 4), but encoding in IT neurons was not perfectly separable (e.g. Figure 3C). Based on this we propose that reading could increase the overall separability of word representations. This was indeed observed recently in the anterior LOC (Agrawal et al., 2019), the homolog of IT cortex in humans. Whether this separability comes with exposure to single letters or to letter strings would be interesting to study. Second, we note that familiarity has widespread effects on IT neurons (Woloszyn and Sheinberg, 2012; Meyer et al., 2014; Srihasam et al., 2014) and on human visual cortex (Agrawal et al., 2019; Gomez et al., 2019). We speculate that reading could lead to enhanced discrimination as letters become familiar, as has been observed recently (Agrawal et al., 2019).

Our results also have important implications for machine vision. They indicate the specific representational rules that need to be embodied in order to solve CAPTCHAs. Indeed, there are deep network design and training principles that implement one or few of the following attributes namely compositionality, sparseness and divisive normalization. An attempt has been made to bring in compositionality (Stone et al., 2017) into deep networks albeit maintaining a veridical mapping between sensory input and internal representation. Randomly deleting connections during training iterations, also called drop-out, as well as non-linearities such as rectified linear units (ReLUs) can bring in sparseness (Hinton et al., 2012; Warde-Farley et al., 2013). Finally, averaging CNN filter responses in a spatial neighborhood (mean-pool) is similar to divisive normalization for broadly tuned neurons and retaining only the highest response in a local neighborhood (max-pool) can approximate aggregation of highly sparse neurons (Nagi et al., 2011). We propose that explicitly encoding separability could lead to vastly improved performance of deep neural networks on distorted letter recognition.

## METHODS

All experiments were performed in accordance with a protocol approved by the Institutional Animal Ethics Committee of the Indian Institute of Science and by the Committee for the Purpose of Control and Supervision of Experiments on Animals, Government of India. Most experimental procedures were common to previously reported studies in our lab (Ratan Murty and Arun, 2015, 2018) and are only briefly summarized below.

### Neurophysiology

We recorded from the left IT cortex of two macaque monkeys (*macaca radiata, Ka* & *Sa*, both male, aged 7 years) using standard neurophysiological procedures. Recording sites were located in the anterior ventral portion of IT. Extracellular wideband signals were recorded at 40 KHz using 24-channel laminar electrodes (Uprobes with 100 µm inter-contact spacing, Plexon Inc). We manually sorted these signals offline into distinct clusters using spike-sorting software (OfflineSorter, Plexon Inc). We selected 139 visually responsive units for further analysis.

### Stimuli

We created a set of 432 unique stimuli. We selected eight letters which included lower- and upper-case letters and numbers (‘a’, ‘d’, ‘g’, ‘s’, ‘T’, ‘y’, ‘7’, ‘K’). Each single letter was rendered at six retinal locations 0.8° apart, three adjacent locations being ipsilateral and three contralateral. Presenting each of the unique letter shapes at six unique spatial locations resulting in a total of 48 possible single letter placements. These locations were chosen to match the letter locations in the longer n-gram strings described in the next section. Letters were rendered in the Times New Roman font and subtended 0.8° visually on an average.

These letters were also used to create 10 bigrams, 10 trigrams all the way to 10 6-grams (Figure S1), resulting in a total of 50 unique n-grams. Thus, there were a total of 98 undistorted single letters and n-grams.

Next we applied three different distortions drawn from commercial captcha algorithms to each of these 98 letter or n-gram placements. In the first, we applied a local fish-eye lens distortion that expands and translates pixels in x and y directions (https://in.mathworks.com/matlabcentral/fileexchange/22573-pinch-and-spherize-filter). We applied a second distortion over and above the first distortion which involved randomly rotations (uniformly chosen from -15 to 15 degrees) of every character in the string. The third distortion was added over and above the second distortion and affected the global shape of the string. It involved bending the entire n-gram about one of two different spline curves. All distortions were implemented using custom scripts written in MATLAB. Since each stimulus was presented in its undistorted form and three distortions, this resulted in a total of 98 x 4 = 392 stimuli.

In addition to these distortions, we also included hard CAPTCHA variants used by Google, Yahoo and other web services (Figure 6). We selected a subset of eight 6-gram strings (‘7Kadgs’, ‘KadgsT’, ‘Ty7Kad’, ‘Tyadgs’, ‘gsTy7K’, ‘sTy7Ka’, ‘y7Kadg’, ‘yadgsT’) which were distorted in four additional ways: (1) rotating the letters in place so that they touched each other; (2) retaining only the letter outlines; (3) adding a fish-eye transformation, rotating letters in place and adding a background line grid and (4) adding a fish-eye transformation, rotating letters in place (uniformly at random between 15 to -15 degrees) and adding a wavy stroke cutting through the letters. These distortions can be seen in Figure 6. These distortions are known to be challenging for computer vision algorithms and are used for human-vs-robot authentication tests (von Ahn et al., 2003) in commercial web services from Yahoo® (touching solid letter and letter outline hard CAPTCHAS) and Google® (hard CAPTCHAS with strike-through and background grid).

### Behavioural task

Each animal was trained to fixate a series of 8 images presented at fixation in return for a juice reward. Each image was shown for 200 ms with an inter-stimulus interval of 200 ms. Thus, each trial lasted 3.2 s. Trials contained either single letters or n-grams, randomly ordered with the constraint that the same stimulus should not repeat in a given trial. The 432 stimuli were organised into 54 blocks of 8 stimuli each and we generated four unique sets of these 54 trials using pseudo-random assignment of letter or n-gram stimuli. During recordings, a set of 54 trials were completed (corresponding to one repetition of the entire stimulus set) before starting another set of 54 trials. N-grams with even number of letters (2-, 4- & 6-grams) were centered on fixation whereas those with odd number of letters (3- & 5-grams) were positioned such that the contralateral side had one extra letter.

### Calculation of response reliability

We obtained a measure of response reliability for each neuron, which represents an upper bound on model performance. To do so, we calculated the average firing rate of the neuron across all stimuli under consideration across the odd-numbered and even-numbered repetitions separately.

### Model fits for letter x location x distortion encoding (Figure 2C-E)

We tested each neuron with 8 letters presented at 6 retinal locations and 4 distortions. We used the firing rate of each neuron (calculated in a 50-200 ms window after stimulus onset) to these 192 stimuli to fit two models. To do so, we first calculated the average shape tuning for the neuron by averaging its response (on odd-numbered repetitions) to every shape across retinal locations and distortions. We proceeded likewise to calculate the average retinal location tuning and average distortion tuning. For the additive model, the response to shape i, location j and distortion k is given by *R*(*s*_*i*_, *p*_*j*_, *d*_*k*_) = *S*_*i*_ + *P*_*j*_ + *D*_*k*_, where *S*_*i*_ is the average shape tuning for shape i, *P*_*j*_ is the average location tuning for location j and *D*_*k*_ is the average distortion tuning for distortion k. Likewise, for the multiplicative model, the response is given by *R*(*s*_*i*_, *p*_*j*_, *d*_*k*_) = *S*_*i*_ ∗ *P*_*j*_ ∗ *D*_*k*_. Both model predictions were scaled and shifted to be in the same range as the observed response. To evaluate the model fit, we calculated the correlation between the model predictions (trained on odd-numbered repetitions) and the firing rate of the neuron on even-numbered repetitions. For Figure 2E, we excluded 37 neurons with high response variability that caused them to have a negative split-half correlation.

### Model fits for n-gram x distortion encoding (Figure 2G-I)

We tested each neuron with 10 n-grams at each of four distortions for each length from 2 to 6 letters. We used the firing rate of each neuron calculated as before to the 40 n-grams of each specific length to fit two models. As before we calculated the average n-gram tuning and average distortion tuning. For the additive model, the response to shape i, distortion k is given by *R*(*s*_*i*_, *d*_*k*_) = *S*_*i*_ + *D*_*k*_, where *S*_*i*_ is the average shape tuning for shape i and *D*_*k*_ is the average distortion tuning for distortion k. Likewise, for the multiplicative model, the response is given by *R*(*s*_*i*_, *d*_*k*_) = *S*_*i*_ ∗ *D*_*k*_. Both model predictions were scaled and shifted to be in the same range as the observed response. As before, we trained the model on the odd-numbered repetitions and calculated its correlation with the firing rate on even-numbered repetitions. For Figure 2I, we excluded 31 neurons with high response variability that caused them to have a negative split-half correlation.

### Model fits for letter combinations (Figure 3)

Each neuron was tested with 10 strings at 4 distortions for each n-gram length (n = 2, 3, 4, 5, 6). We used the firing rate of each neuron calculated as before to the 40 n-grams of a given n-gram length to fit a model based on single letters. For each n-gram (e.g. “yadgsT”), we compiled the corresponding predicted single letter responses from the multiplicative model (shape x distortion x location) (e.g. to “y” at location 1, “a” at location 2, etc). Using the raw firing rates to single letters yielded qualitatively similar results. We compiled the n-gram responses (for each length) and the corresponding single letter responses into the matrices **y** and **X** respectively where **y** is a 40 x 1 vector containing the responses to the 40 n-grams of that length in odd reps, **X** is a 40 x n matrix containing the corresponding single letter responses at each of n retinal locations together with a constant term. We then modelled the full set of 200 unique n-gram responses as a single equation **z = Ab**, where **z** is a concatenation of all **y** n-gram response vectors at each length (n = 2, 3, 4, 5, 6) and the letter response matrix **A** was a 200 x 25 matrix in in which corresponding single letter responses at each length were entered into a separate set of columns. In other words, the matrix **A** had a total of 25 columns made up of 3 columns for bigram letter responses (two weights & one constant), 4 columns for trigrams (three weights and one constant), and so on. The regression weight vector b is a vector of 25 unknowns representing the weights of each location within the n-gram for each particular n-gram length. To avoid overfitting since these models have relatively few parameters compared to observations, model predictions were calculated using cross-validation: we trained the model on 80% of the data (i.e. 160 stimuli) and calculated its prediction on the held-out 40 stimuli. These predictions were compared with the observed responses for the plots in Figure 3B & 3C. For Figure 3D, since the goal was to analyse the summation weights, we fit the model only once on the full set of 40 n-gram responses and used these weights for averaging. In addition, neural responses were normalized by the maximum firing rate across all 40 stimuli, so that the resulting weights could be averaged across neurons without being unduly influenced by high firing neurons. For Figure 3C, we excluded 48 neurons with high response variability that caused them to have a negative split-half correlation.

### Decoding character identity for every location in a n-gram

For each neural population, we trained linear classifiers to decode the individual letters at each retinal location. To simultaneously decode all n-grams, we included a blank as a character so that there were 9 possible labels (8 letters + 1 blank) at each retinal location. For every retinal location, a n-gram response was assigned the label corresponding to the character contained at that location. i.e.; the n-gram ‘adgsT’ would have the letter labels {‘blank’, ‘a’, ‘d’, ‘g’, ‘s’, ‘T’} in retinal locations 1 through 6. The number of labeled training instances for each label class were balanced to avoid classifier bias. In this manner, we trained six 9-way linear classifiers for each of the six retinal locations. Decoding accuracy is reported throughout as the average performance across these six retinal locations. Since there are 9 classes in the classifier, chance performance is 1/9 = 11%. Since blank spaces occurred more frequently at eccentric retinal locations, we calculated the average number of observations across the 8 character classes at a given retinal location and sampled an equal number randomly from the set of responses corresponding to blank spaces. We did this to prevent the classifier from learning trivial decision boundaries such as classifying all test observations as blanks.

### Creation of artificial neurons with separable responses (Figure 4A)

We created a population of 1,000 artificial neurons with separable responses as follows. Each neuron was initialized with tuning for letter, location and distortion that could be sharp or broad. For broad tuning, we selected responses to be uniformly sampled from 0 to 1 generator (rand function in MATLAB). For sharp tuning, we chose only one response to equal 1 and the remaining to be 0 according to a sparse matrix generator (sprand function in MATLAB, with density 1/(n-gram length)). For mixed tuning, we created neurons that were selective from only one or more letters but not to the remaining ones (sprand function in MATLAB, with density 1/k, k = 1, 2, …, 6). A given neuron was chosen as sharply (or broadly) tuned for all three attributes (shape, location, distortion) since IT neurons are known to have this property (Zhivago and Arun, 2016). For the mixed population of neurons, each neuron was randomly assigned (with equal probability) to be sharply or broadly tuned. We then calculated the response of each neuron to distorted letters and strings by applying either additive or multiplicative. Finally, the response to n-grams was calculated as the sum of the responses to single letters at the corresponding retinal locations and divided by the number of letters (if divisive normalization was selected). In this manner, we created several groups of artificial neurons: broad tuning with multiplicative/additive integration with/without normalization, sharp tuning with multiplicative/additive integration with/without normalization and mixed tuning with multiplicative/additive integration with/without normalization.

To truly test the capabilities of CAPTCHA decoding for each synthetic neuron population, we calculated the response of each group to 17,920 n-gram stimuli. We created this multi-letter n-gram dataset from the same 8 characters we used in our original set of stimuli. For n-gram lengths varying from 2 to 6, we first assigned characters without replacement to the same adjacent retinal locations that were used in our original set of n-grams (e.g.; central-most two locations for bigrams), non-letter positions were filled with blanks. This procedure yielded 8×8=64 unique strings for bigrams (eg; **aa**, **ab** where * denotes blanks). In this manner we obtained 64, 512, 4096, 32768 and 262144 strings of lengths 2 to 6 respectively. We then performed circular-shifts on these strings to obtain strings with new spatial mappings for adjacent characters for example the string **ab** gave the unique strings ***ab*, ****ab, b****a and ab**** using this procedure. We then created more unique strings by letters across retinal locations obtained more strings such as *a**b*, *a***b and included in the dataset if they were not already present. At this stage we obtained 28736, 3584, 4416, 32768, 262144 strings of lengths 2 to 6 respectively. To avoid training biases, we sampled from the longer string sets to obtain a dataset with 17,920 unique strings with 3,584 n-grams of every length.

### Separability in deep convolutional neural networks

We analyzed four deep convolutional neural networks for their responses to our stimulus set. The first network served as a baseline and is a VGG-16 architecture trained for object classification on the ImageNet dataset (Russakovsky et al., 2015). The VGG-16 model we used had 5 convolution stages that had 3×3 filters and 64, 128, 256, 512, 512 channels respectively and each convolution was accompanied by 2×2 max-pooling and RelU (Rectified linear unit) operations (Simonyan and Zisserman, 2014).

We also selected three deep networks trained for text decoding using images as inputs (Jaderberg et al., 2014, 2016). These networks were trained in these studies using a large-scale dataset created by generating words of lengths up to 23 characters and rendering them using a variety of distortions, fonts, border/shadow rendering, noise and background information statistics. These three models have a common backbone architecture consisting of input (32×100×1), convolutional (32×100×64, 16×50×128, 8×25×256, 4×13×512) and fully connected (1×1×4096) stages, before being connected to either 23 output stages (each 1×1×37) for *charnet*, a single n-gram output stage (1×1×10,000) for *ngramnet* or a single word output stage (1×1×90,000) for *dictnet*.

The first network, which we denote as *charnet*, was trained to correctly identity the letter or number at each retinal location in words of up to length 23. Words were left aligned in a large-scale training dataset of approximately 8 million words. The final decision layer for *charnet* comprised of 851 nodes corresponding to character identities (26 letters + 10 numbers + 1 blank space) for each of 23 locations. The second network, which we denote as *ngramnet*, was trained to detect the presence of every n-grams present in a word out of a possible 10,000 n-grams. For example, when the input image contains the string ‘ad’, this network will return the n-grams {‘a’, ‘d’, ‘ad’}. The third network, which we denote as *dictnet*, was trained to identify whole words from a dictionary of 90,000 possible words.

Because information in all four networks passes through a fully connected stage with 4096 nodes before the final classification stage, we selected this layer for our analyses. To analyze the separability of distorted letter strings in each network, we generated its activations for the stimuli in our neural recordings, and performed the exact same analysis as for real neurons.

## ACKNOWLEDGEMENTS

This work was supported by a DST CSRI post-doctoral fellowship (HK) from the Department of Science and Technology, Government of India and by Intermediate and Senior Fellowships (500027/Z/09/Z and IA/S/17/1/503081) from the Wellcome Trust-DBT India Alliance to SPA.

## Supplementary material for

### Supplementary Figures

**Figure S1.**
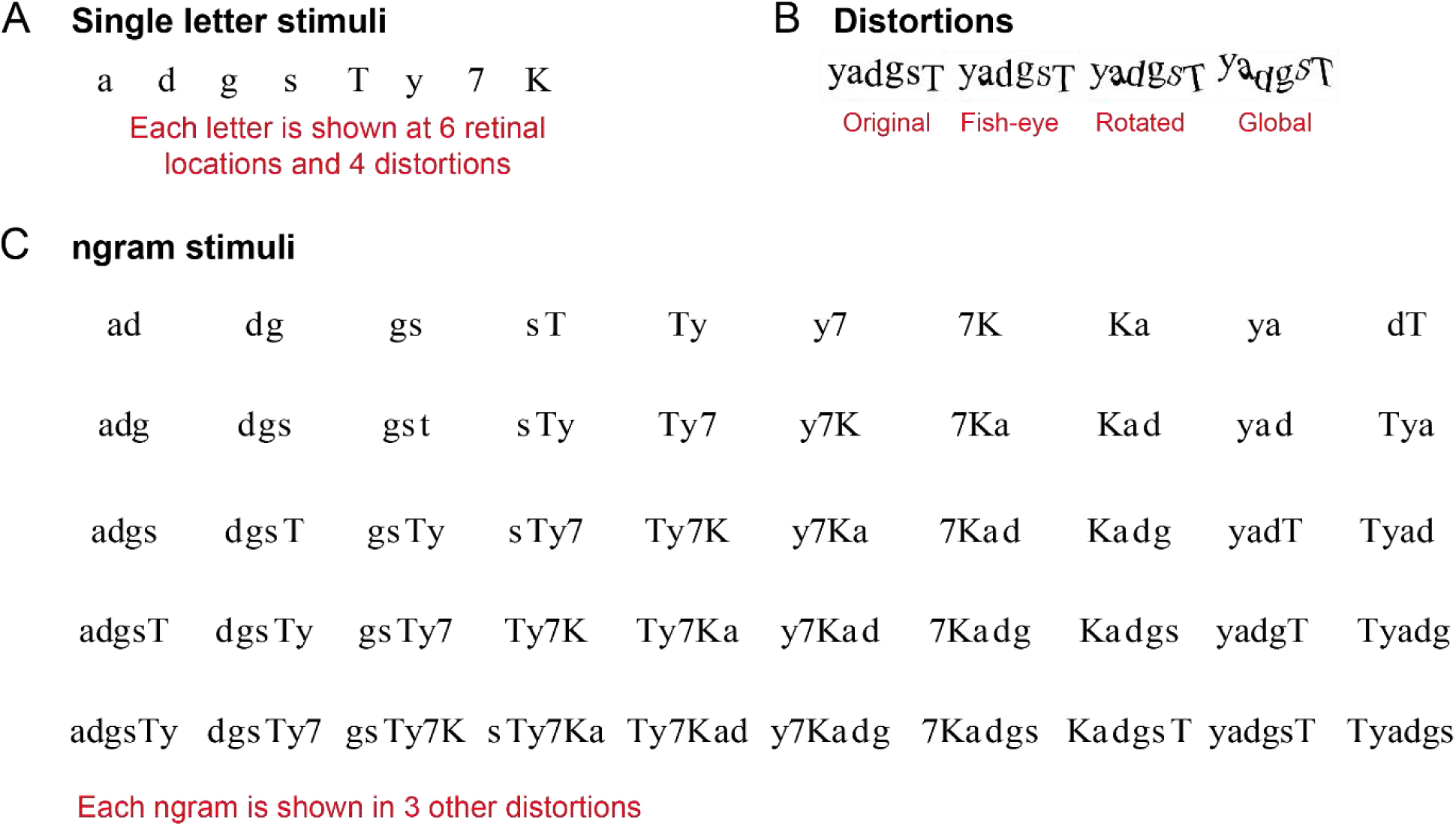
Stimuli used in neural recordings. (A) Eight single letters were presented at 6 retinal locations and 4 distortions. (B) Each stimulus was distorted in three ways: fish-eye, rotated and global. (C) Ngram stimuli were created by combining the 8 single letters in various ways, and ensuring that longer ngrams contained several smaller ngrams.

**Figure S2:**
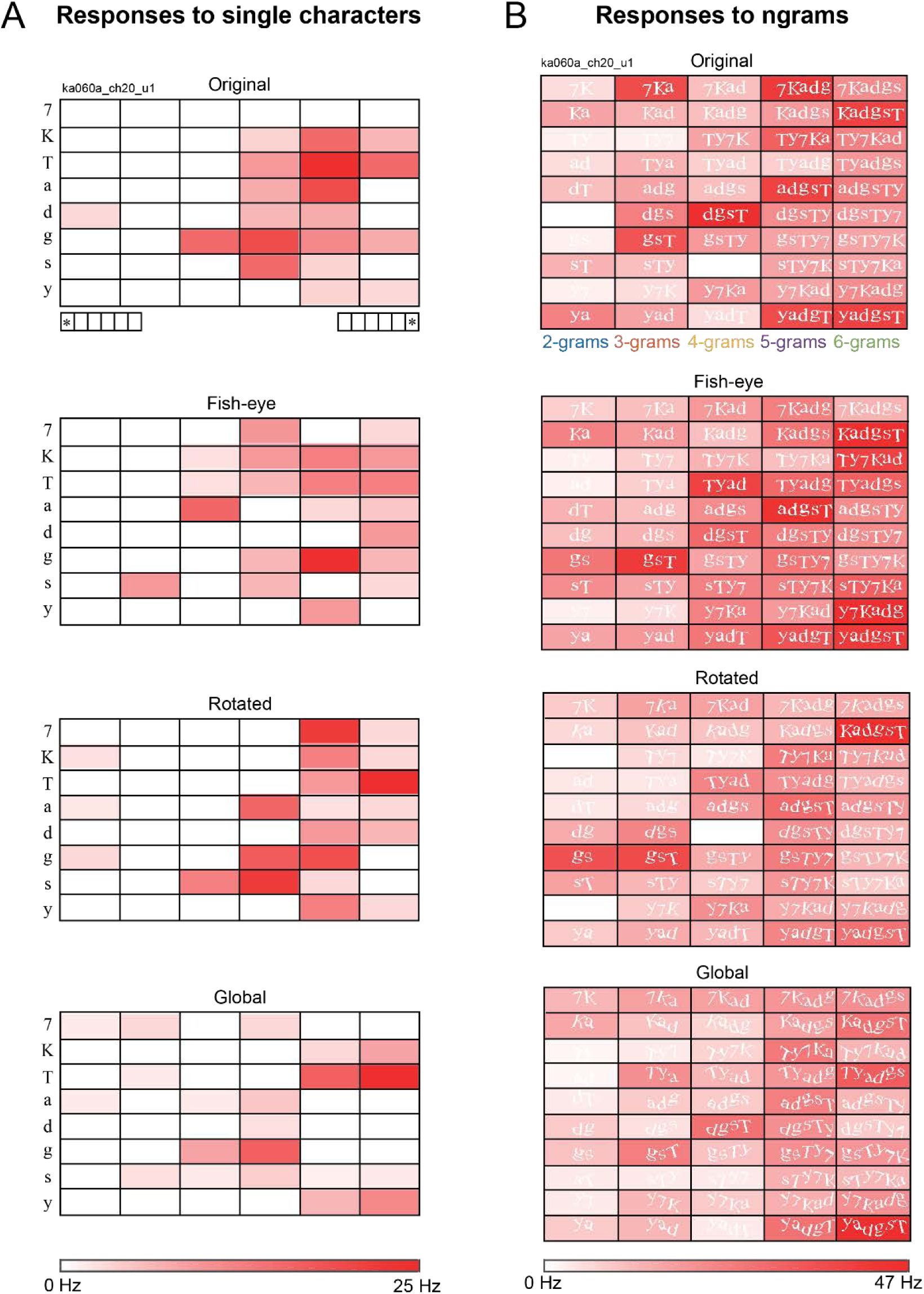
Responses of the neuron ka60_ch20_u1 to all stimuli. (A) Responses to single characters that were shown at 6 retinal locations and 4 different distortion versions. Responses have been grouped by the distortion type. (B) Responses to ngrams that were shown at fixation in one of 4 different distortion versions. Responses have been grouped by the distortion type.

**Figure S3.**
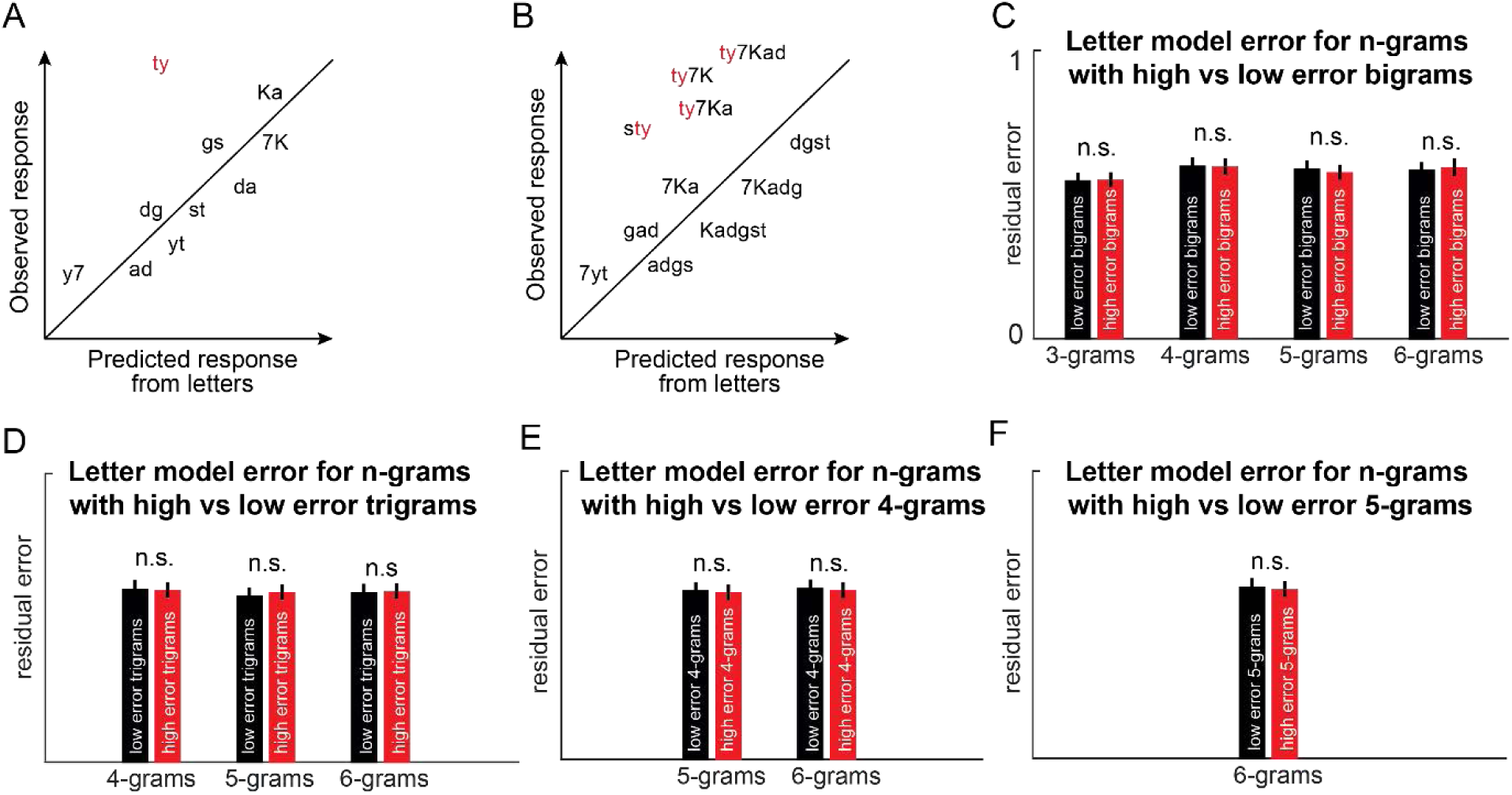
Error analysis for combination detectors. (A) Consider a neuron selective for the bigram “ty”– by definition it would mean that its response to “ty” systematically deviates from the prediction from single letters. (B) If this neuron is consistently detecting the bigram “ty” its response to any longer string containing “ty” should deviate in the same direction. (C) To assess this possibility, we sorted bigrams according to the bigram model error into two groups, one with higher model errors and the other with lower model errors. We then calculated the average model error for longer strings containing high-error bigrams and low-error bigrams after fitting the linear model to strings of each length. There were no systematic differences in model error as would be expected from combination detectors. (D) Similar analysis for trigram detectors. We sorted all longer strings (4-grams, 5-grams and 6-grams) into those containing high-error trigrams and low-error trigrams. We observed no systematic differences in model error as would be expected from trigram detectors. (E) Same as (D) but for 4-gram detectors. (F) Same as (D) but for 5-gram detectors.

**Figure S4.**
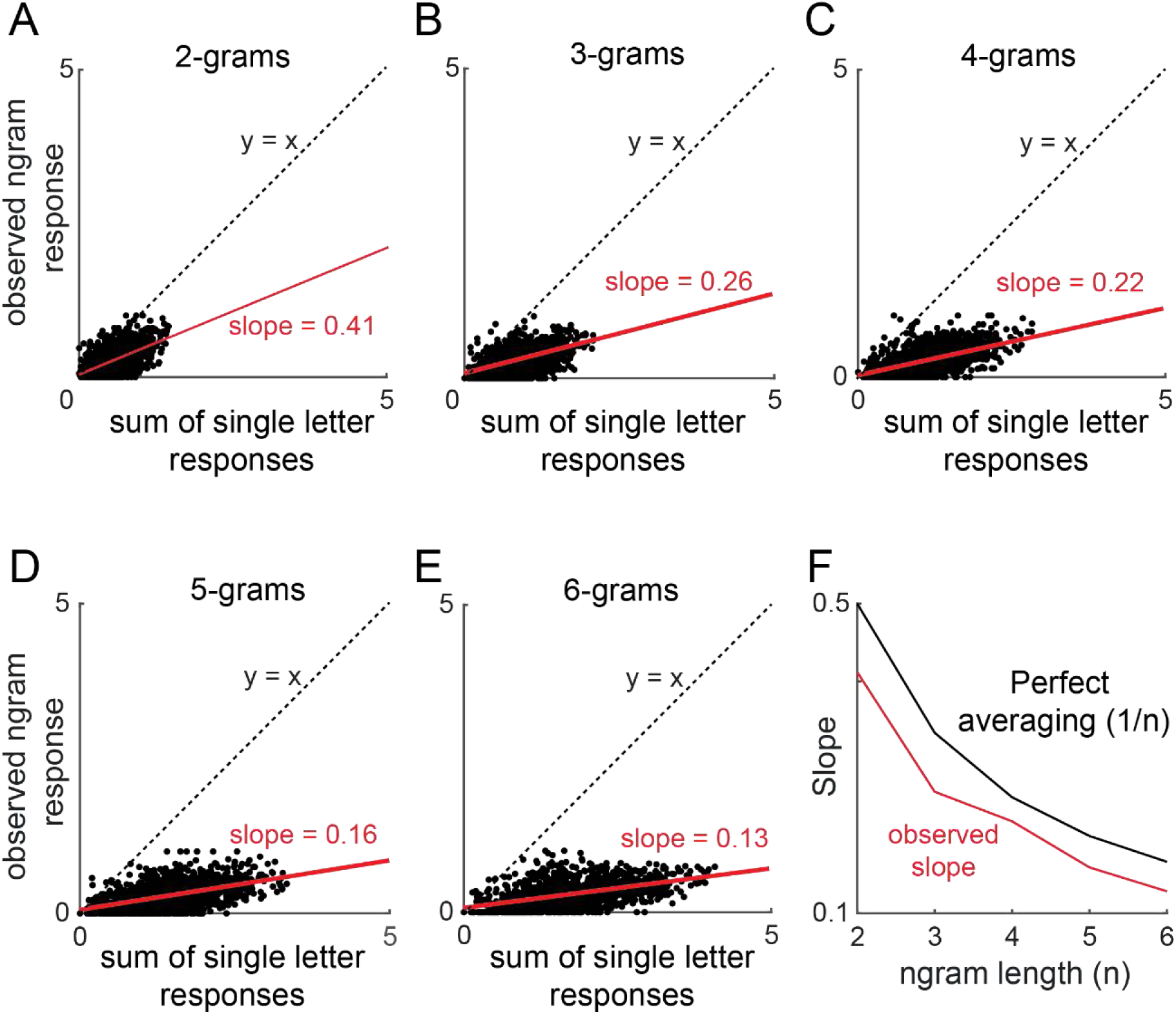
Divisive normalization in ngram responses. (A) Normalization weights calculated as the slope of best fit line that predicts observed ngram responses at each length using the sum of observed single character responses at corresponding retinal locations. Only undistorted strings and characters have been used and neuronal firing rates have been normalized for every neuron. Each plot has 1390 points corresponding to 10 unique ngrams at each ngram length, from every neuron. (B) Estimated slopes of the best fit line are plotted against ngram length.

